# Growth orientations, rather than heterogeneous growth rates, dominate jaw joint morphogenesis in the larval zebrafish

**DOI:** 10.1101/2022.01.13.476166

**Authors:** Josepha Godivier, Elizabeth A. Lawrence, Mengdi Wang, Chrissy L. Hammond, Niamh C. Nowlan

## Abstract

In early limb embryogenesis, synovial joints acquire specific shapes which determine joint motion and function. The process by which the opposing cartilaginous joint surfaces are moulded into reciprocal and interlocking shapes, called joint morphogenesis, is one of the least understood aspect of joint formation and the cell-level dynamics underlying it are yet to be unravelled. In this research, we quantified key cellular dynamics involved in growth and morphogenesis of the zebrafish jaw joint and synthesised them in a predictive computational simulation of joint development. Cells in larval zebrafish jaw joints labelled with cartilage markers were tracked over a forty-eight hour time window using confocal imaging. Changes in distance and angle between adjacent cell centroids resulting from cell rearrangement, volume expansion and extracellular matrix (ECM) deposition were measured and used to calculate the rate and direction of local tissue deformations. We observed spatially and temporally heterogeneous growth patterns with marked anisotropy over the developmental period assessed. There was notably elevated growth at the level of the retroarticular process of the Meckel’s cartilage, a feature known to undergo pronounced shape changes during zebrafish development. Analysis of cell dynamics indicated a dominant role for cell volume expansion in growth, with minor influences from ECM volume increases and cell intercalation. Cell proliferation in the joint was minimal over the timeframe of interest. Synthesising the dynamic cell data into a finite element model of jaw joint development resulted in accurate shape predictions. Our biofidelic computational simulation demonstrated that zebrafish jaw joint growth can be reasonably approximated based on cell positional information over time, where cell positional information derives mainly from cell orientation and cell volume expansion. By modifying the input parameters of the simulation, we were able to assess the relative contributions of heterogeneous growth rates and of growth orientation. The use of uniform rather than heterogeneous growth rates only minorly impacted the shape predictions whereas isotropic growth fields resulted in altered shape predictions. The simulation results suggest that growth anisotropy is the dominant influence on joint growth and morphogenesis. This study addresses the gap of the cellular processes underlying joint morphogenesis, with implications for understanding the aetiology of developmental joint disorders such as developmental dysplasia of the hip and arthrogryposis.

## Introduction

Synovial joints are complex structures connecting skeletal elements while allowing different types of motion. In early limb embryogenesis, the cartilaginous anlagen of the future skeletal elements are initially uninterrupted (Yang 2013). A zone of compact and interconnected cells, called the interzone, emerges marking the location of the future joint. Physical separation of the skeletal elements occurs by cavitation of the interzone while the two opposing surfaces mould into reciprocal and interlocking shapes in a process known as joint morphogenesis (Pacifici et al. 2005, Chijimatsu and Saito 2019, Rux et al. 2019). A variety of distinct and complex joint shapes, which are specific to anatomical sites and allow distinct motions, emerge from this process; examples of joint diversity are the hinge joint of the knee and the ball and socket of the hip. This process by which joints acquire their shapes has important ramifications for joint health and function. For example, sub-optimal hip joint shape is believed to be a key risk factor in early onset osteoarthritis (Sandell 2012, Faber et al. 2020). However, the mechanisms underlying the emergence of joint shape remain poorly understood.

A small number of studies have identified cell activities involved in joint growth and morphogenesis. Work on embryonic murine synovial joints have shown that a continuous influx of pro-chondrogenic cells contributes to joint morphogenesis (Shwartz et al. 2016), with evidence that asymmetric influx and proliferation of these cells enables the emergence of asymmetric shape features (Zhang et al. 2020). The maintenance of cell fate has also been shown to be essential for joint cavitation and morphogenesis. Absent muscle contraction results in premature differentiation of joint pro-chondrocytes with consequences for joint shape in the embryonic murine elbow (Kahn et al. 2009). The roles of cell size, orientation and intercalation in developing zebrafish and murine joints have been identified (Shwartz et al. 2012, Brunt et al. 2015) and differential cell volume expansion and cell rearrangements were shown to be key factors for thickening and organisation in postnatal murine articular cartilage (Decker et al. 2017). Cell proliferation and cell death do not majorly impact morphogenesis in postnatal murine articular cartilage (Decker et al. 2017). These observations provide insights on the cellular dynamics underlying joint morphogenesis, but there is a lack of understanding of the contribution of each of these processes to joint growth and morphogenesis. The research question we tackle in this paper is how a complex range of dynamic cellular activities combine to enable the formation of specific shape features in synovial joints.

Computational models enable the synthesis of experimental data and a means to test hypotheses via simulation. In previous work from our group, (Giorgi et al. 2014), the emergence of different joint shapes based on types of simulated fetal movements was predicted in a mechanobiological simulation. A simulation of hip joint development revealed how asymmetric movements can result in altered shapes resembling those seen in developmental dysplasia of the hip (Giorgi et al. 2015). A later simulation using aspects of the same model from another group investigated the impact of muscle mass and anatomy on development of the glenohumeral joint and was able to predict the formation of brachial plexus birth injury (Dixit et al. 2020). The limitation of most simulations of joint morphogenesis is that they are based on simplified or extrapolated cell activities. Our simulations and those of others (Shefelbine and Carter 2004, Giorgi et al. 2014, Dixit et al. 2020) have modelled the biological contribution to growth as being proportional to chondrocyte density, based on a study by Heegaard et al. (1999), in which chondrocyte density was approximated based on the grey level distribution of a section of a human interphalangeal joint. As cellular processes orchestrate any changes in joint shape, the lack of a more precise and specific characterisation of cell-level activities to joint growth and morphogenesis is a striking gap. Simulations of joint growth and morphogenesis based upon accurately tracked cell activities will provide insights into the mechanisms underlying prenatal joint growth and morphogenesis.

There is a growing body of research on quantifying cellular dynamics involved in growth and morphogenesis using computational tools. Rubin et al. (2021) built 3D maps of cell morphologies from light-sheet images of the embryonic murine tibia. Cell density, surface area, volume and orientation were quantified and spatially analysed revealing that differential cell volume expansion underlies tissue morphogenesis of the developing growth plates. Stern et al. (2021) quantified cell dynamic behaviours, such as proliferation and intercalation, in the epithelial sheet of the Drosophilia embryo and evaluated their impact on gastrulation in terms of area expansion and tissue stretching. Heller et al. (2016) developed an automated image analysis toolkit for epithelial tissues called EpiTools which enables spatial and temporal morphometric analysis of time lapse images taken at high temporal and cellular resolution—namely cell surface area, shape, division, orientation and intercalation. Applied to Drosophilia wing imaginal disc, this toolkit provided new understanding of the role of cell rearrangements underlying tissue growth and morphogenesis. Others have been able to directly quantify tissue growth based on cell level data using lineage tracing (Marcon et al. 2011, Morishita et al. 2015, Suzuki and Morishita 2017, Tozluoglu et al. 2019). Quantitative maps of tissue deformation coupling growth rates and anisotropy were obtained in developing chick limbs (Marcon et al. 2011, Morishita et al. 2015, Suzuki and Morishita 2017) and in the Drosophilia wing disc (Tozluoglu et al. 2019). These studies showed that spatially and temporally heterogenous growth patterns as well as growth anisotropy are key drivers of morphogenesis, while uniform growth rates do not lead to correct shape predictions. We are not aware of any similar studies quantifying the cellular dynamics of joint morphogenesis. Such characterisation combined with computational simulation of joint growth will help us to unravel different contributions to joint morphogenesis, including the roles of cell volume changes and rearrangements as previously highlighted in other growing tissues.

In this research, we quantify the cell-level dynamics during joint morphogenesis by tracking cell activities in high resolution in larval zebrafish jaws, then synthesise them in a predictive computational simulation of joint development. We use the simulation to test if growth heterogeneity or growth orientation are the dominant influences on joint growth and morphogenesis. This paper addresses the gap in knowledge on the cellular processes and dynamics leading to morphogenesis of developing joints.

## Methods

### Zebrafish husbandry/Zebrafish lines

Fish were maintained as described in Aleström (2020). All experiments were approved by the local ethics committee (Bristol AWERB) and performed under a UK Home Office Project Licence. Transgenic lines *Tg(col2a1aBAC:mCherry)* (Mitchell et al. 2013) and *Tg(−4.9sox10:eGFP)* (Carney et al. 2006) have been previously described.

#### CHARACTERISING GROWTH FROM CELL-LEVEL DATA IN ZEBRAFISH JAW JOINTS

##### Zebrafish jaw joint live imaging

Ten jaw joints from double transgenic *Tg(col2a1aBAC:mCherry; −4.9sox10:eGFP)* larvae were imaged at 12-hour intervals from 3.5 to 5.5 days post fertilisation (dpf) using a Leica SP8 confocal microscope with a temperature-controlled chamber set to 28°C. Images centred on the joint line, as marked by a red box in Figure 1A, were acquired with a 20× HCX PL APO lens at a resolution of 512 × 512 px. Prior to imaging, larvae were anaesthetised in 0.1 mg ml-1 tricaine methanesulphonate (MS222) and mounted in a ventral orientation in warm 1% low melting point (LMP) agarose. Following imaging, the larvae were flushed from the agarose using Danieau’s buffer and kept in separate wells of a 24-well plate between imaging timepoints.

**Figure 1:**
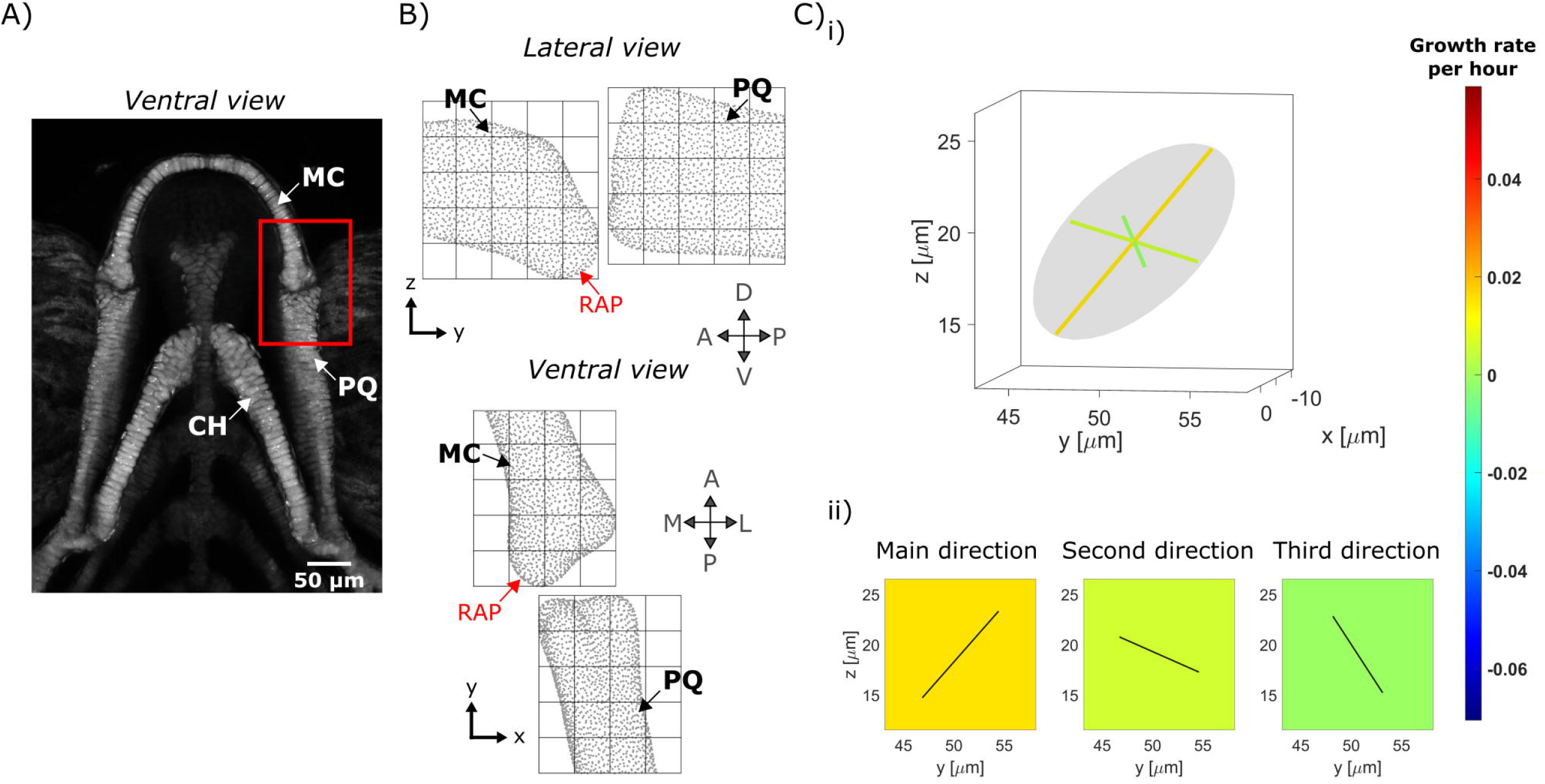
Growth map calculations in larval zebrafish jaw joint. A) Maximum projection of ventral confocal image stacks of the jaw from a larval zebrafish aged 5 dpf expressing *Tg(Col2a1aBAC:mcherry)* cartilage marker; red box shows the jaw joint for which morphogenesis is characterised in this study; B) A grid marks out the regions (ROIs) of the anterior MC and posterior PQ joint elements in which growth is characterised. Each cube side length is 15μm. C) i) The growth rate calculated for each ROI is represented by an ellipsoid with orthogonal axes. ii) The ellipsoid’s radii and the orientation of its axes are used to generate a growth map for each of the ellipsoid’s radii in the lateral plane; growth rate is represented by the square’s colour while the direction of growth is shown by solid black lines in the corresponding square. MC: Meckel’s cartilage, PQ: Palatoquadrate, CH: ceratohyal, RAP: retroarticular process, A: Anterior, P: Posterior, L: Lateral, M: Medial, D: Dorsal, V: Ventral.

##### Cell segmentation and tracking

Consecutive image stacks with *sox10:eGFP* chondrocyte marker were filtered in Fiji (Schindelin et al. 2012). 3D Fast Filters-OpenGray, 3D Edge and Symmetry and 3D Morphological filters from the 3D ImageJ Suite plugin (Ollion et al. 2013) were applied in this order with the parameters supplied in Table 1. Once filtered, morphological segmentation followed by Inertia Ellipsoid filtering using Fiji’s MorpholibJ plugin (Legland et al. 2016) were performed to extract the 3D cell centroids’ coordinates in the joint at each timepoint. Segmentation results were then cleaned and used to manually track joint cells between images from two consecutive timepoints using manual labelling in MATLAB (R2018a, The MathWorks, Inc., Natick, Massachusetts, United States). Cells in which *col2a1aBAC:mCherry* cartilage marker expression was absent were considered part of the interzone and not tracked. Due to image resolution and segmentation quality some image stacks were discarded from the analysis, and the final sample numbers per timepoint are detailed in Figure 4. At each timepoint, cells in the joint were counted to assess proliferation, and the volume occupied by the tracked cells was calculated to assess cell volume expansion.

**Figure 2:**
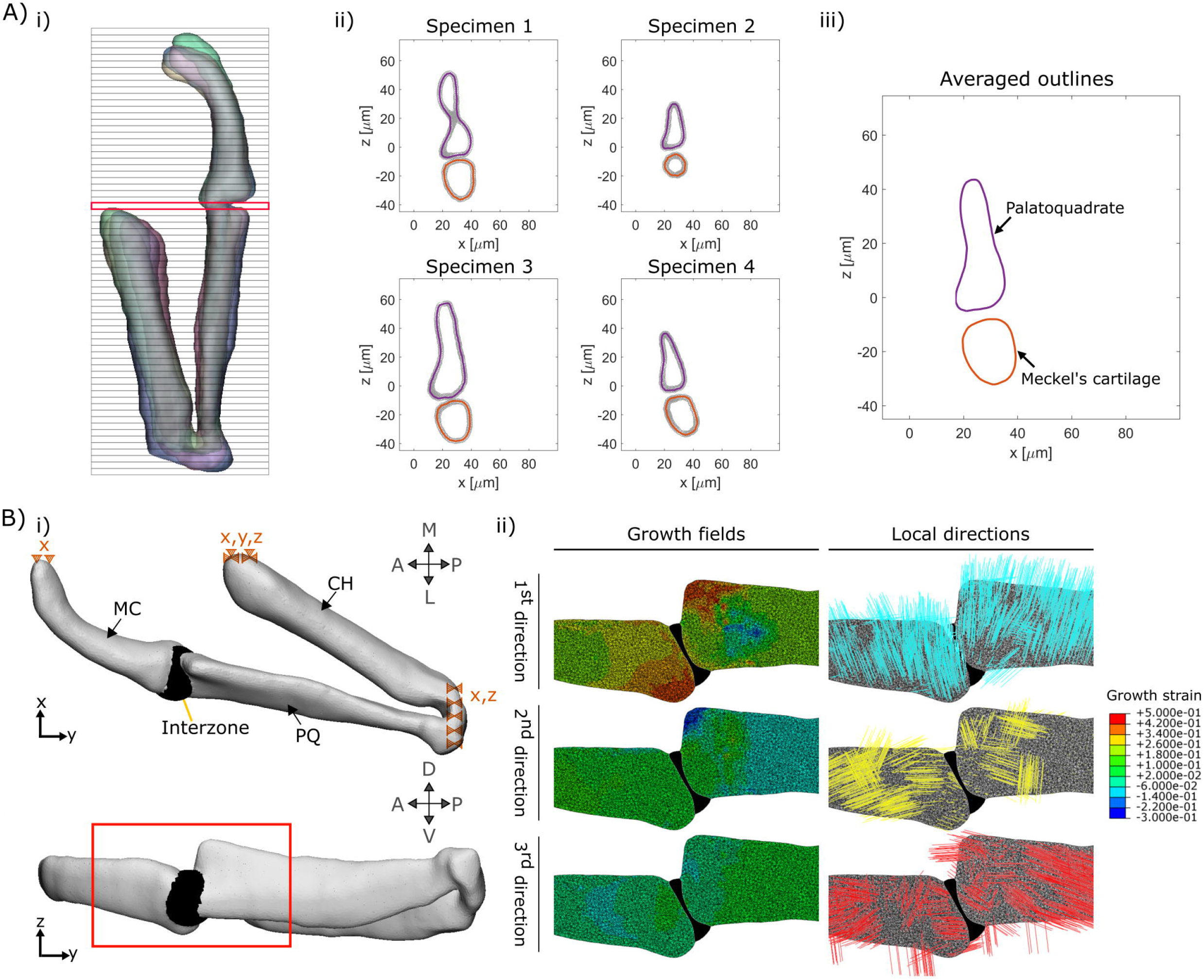
Integration of the growth maps in a finite element model. A) The first step in constructing our FE model is to obtain an average geometry for each timepoint (3.5, 4, 4.5, 5 and 5.5 dpf). For each timepoint, half jaw shapes are aligned and sliced transversally (i). For each slice, the shape outlines of each sample (four here) are obtained (ii) then averaged (iii). The slide marked in red in (i) is shown as an example in (ii) and (iii). B) i) An FE model is generated based on the averaged shape outlines; the joint interzone is added and the areas marked with dashed triangles are constrained in the specified dimensions (e.g., x). ii) Section of the joint in the lateral plane showing the growth fields which are applied to the model along with their associated directions. The view is marked by a red box in (i). MC: Meckel’s cartilage, PQ: Palatoquadrate, CH: ceratohyal, A: Anterior, P: Posterior, L: Lateral, M: Medial, D: Dorsal, V: Ventral.

**Figure 3:**
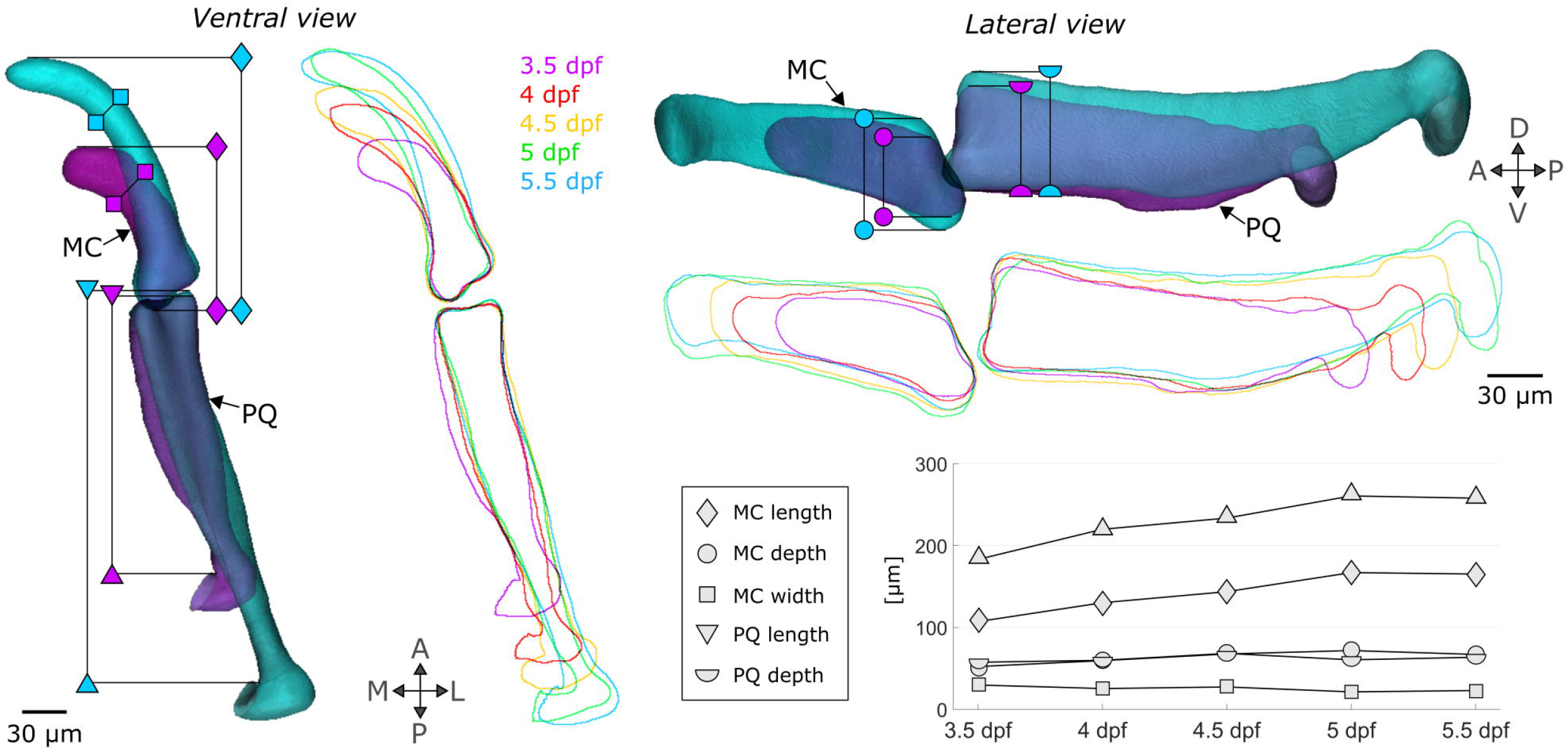
Shape changes between 3.5 and 5.5 dpf in zebrafish jaws. Superimposed 3.5 (purple) and 5.5 (turquoise) dpf 3D average shapes and 3.5 to 5.5 dpf average shape outlines in the ventral and lateral planes. The shape features which are observed to change as the jaw develops are marked with specific symbols (diamond: MC length, square: MC width, circle: MC depth, triangle: PQ length, semi-circle: PQ depth). MC: Meckel’s cartilage, PQ: palatoquadrate, A: Anterior, P: Posterior, L: Lateral, M: Medial, D: Dorsal, V: Ventral.

**Figure 4:**
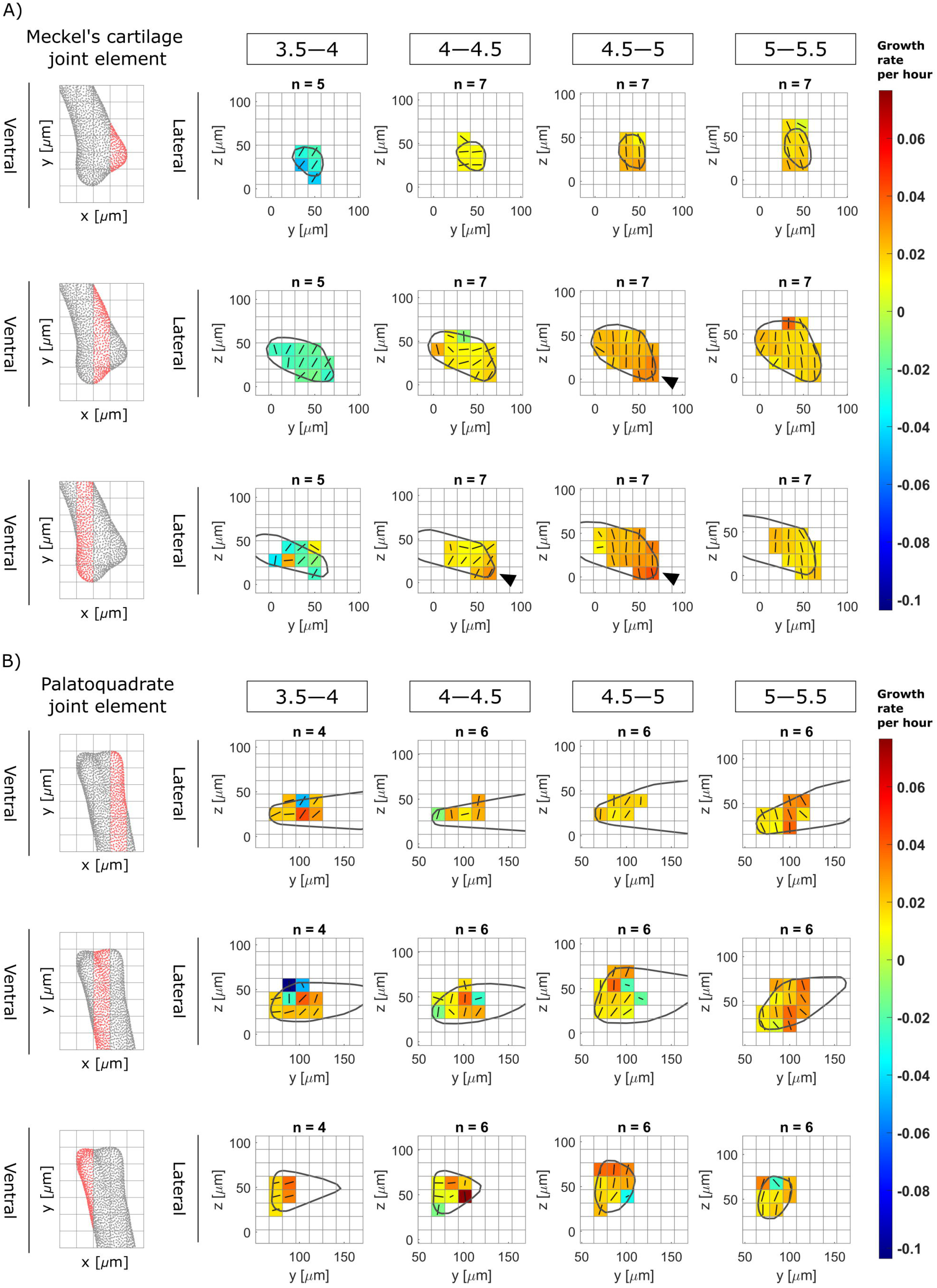
Growth rates from 3.5 to 5.5 dpf in zebrafish jaw joint exhibits spatial and temporal patterns. Maps showing growth rates along the main direction for growth (major axis of the ellipsoid) and their associated directions for each time window (3.5—4, 4—4.5, 4.5—5 and 5—5.5) in the anterior Meckel’s cartilage (A) and posterior Palatoquadrate (B) joint elements in the lateral plane. Growth rates are represented by colours while the direction is shown by solid black lines. Results are displayed across the rudiment’s width; views in the ventral plane of each section are displayed on the left panels. Black arrows in (A) show areas of elevated growth rates.

**Table 1:** Filters applied to larval zebrafish jaw joint image stacks before cell segmentation.

| Filters | Parameters |
| --- | --- |
| 3D Fast Filters-OpenGray | Isotropic radius: 2 pixels |
| 3D Edge and Symmetry | Canny: 0.6 |
| 3D Morphological Filter | Operation: closing<br>Element shape: diamond<br>Isotropic radius: 2 pixels |

##### Growth maps calculations

For each 12-hour interval time window (3.5—4, 4—4.5, 4.5—5 and 5—5.5), joint shapes were extracted from the consecutive image stacks with *col2a1:mcherry* chondrocyte marker in Mimics (Materialise NV, Leuven, Belgium) and aligned in 3-matic (Materialise NV, Leuven, Belgium). Any transformation which was applied to the joint shapes in 3-matic was consistently applied to the corresponding centroids in MATLAB. A cubic grid of side length fifteen microns was superimposed on the aligned joints to divide them into regions of interest (ROIs) as shown in Figure 1B. For each ROI, cells within the ROI’s limits were detected and their adjoining cells were listed. Vectors linking the centroids of adjacent cells were created. In each of the ROIs, a “statistical velocity gradient” was calculated based on vector length and orientation variations between images from consecutive timepoints using the method described by Graner et al. (2008). This gradient quantifies local distortions and rearrangements, such that if cells within an ROI grow or intercalate, or if extracellular matrix is built, the distance between cell centroids, and therefore the geometry of the tissue, change. The statistical velocity gradient can be represented by an ellipsoid with orthogonal axes, as illustrated in Figure 1C. The orientation of the axes and their associated radii respectively correspond to the direction and rate of local tissue geometry deformation. Maps of local strain rates with the associated directions of deformation were generated from each of the three ellipsoid’s axes as shown in Figure 1C. These maps are referred to hereafter as growth maps. For each time window, growth maps were calculated for each of the samples and then averaged. Within each ROI, strain rates that lay outside the interquartile range were removed from the averaging.

Raw, filtered, and segmented confocal image stacks, along with MATLAB codes for cell tracking and growth rate calculations are available at doi: 10.5281/zenodo.5769854.

#### SIMULATING ZEBRAFISH JAW GROWTH WITH A FINITE ELEMENT MODEL

##### Shape generation

Confocal image stacks of four to five larval zebrafish jaws (encapsulating the Meckel’s cartilage, the palatoquadrate and the ceratohyal, see Figure 1A) from the transgenic line *Tg(col2a1aBAC:mCherry)* were taken with a Leica SP8 confocal microscope at the “endpoints” of each time window (3.5, 4, 4.5, 5 and 5.5 dpf) using the methodology described above. A 3D Gaussian grey filter with isotropic radius 3.0 pixels was applied to the image stacks in Fiji. These were imported in Mimics to be segmented and the resulting 3D surfaces were aligned. Only half-jaws (separated at the level of the midsagittal plane) were segmented, as shown in Figure 2A(i). The half-jaws were imported into MATLAB and were divided into slices in the transversal plane as shown in Figure 2A(i). For each slice, a shape outline was obtained for each specimen from the shape vertices and an average outline was generated as shown in Figure 2A(ii-iii). Averaged shape outlines were saved as image stacks and imported into Mimics where the resultant average half-jaw shape was generated. Also in Mimics, the interzone was added as a volume filling the gap between the two joint elements using Boolean operations, with the interzone’s external boundaries approximated based on imaging data (Brunt et al. 2016). Finally, a non-manifold assembly combining the half-jaw and the interzone was generated as shown in Figure 2B(i). In 3-matic, the non-manifold assembly was meshed with ten node tetrahedral elements and exported to Abaqus CAE (Dassault Systemes, 2019) where a model for each 12-hour time window was created.

##### Material properties and boundary conditions

All cartilaginous regions (Meckel’s cartilage (MC), palatoquadrate (PQ) and ceratohyal) were assigned homogeneous isotropic elastic material properties with Poisson’s ratio 0.3 and Young’s Modulus (YM) 54.8 kPa based on nanoindentation measurements taken on 5 dpf wild type zebrafish jaw joints (Lawrence et al. 2021). The interzone was assigned isotropic elastic material properties with Poisson’s ratio 0.3 and YM set at 0.25% of the cartilaginous YM based on Brunt et al.’s original study where this ratio between the two YM was found to facilitate physiological jaw displacements when muscle loading was applied (Brunt et al. 2015). The ceratohyal does not form part of the region of the jaw joint of interest (see Figure 1), but was needed for coherent boundary conditions. The following boundary conditions were applied, as illustrated in Figure 2B(i): the anterior end of the ceratohyal was fixed in all directions, only anteroposterior translations of the posterior end of the ceratohyal were allowed and translations of the anterior end of the Meckel’s cartilage in the lateromedial direction were prevented.

##### Growth maps integration

For each 12-hour period, strains derived from the growth maps were imported into Abaqus CAE as three distinct analytical mapped fields and applied to the model. The coordinates of the ROI centres were assigned the calculated strains and interpolation was performed between ROI centres to assign strains to each element lying within the ROIs’ limits. Local material orientations matching the local directions for growth were assigned to the joint elements. Elements whose nodes’ coordinates were contained within an ROI were all assigned the directions for growth of this ROI. Direction 1 is the main direction for growth (corresponding to the major axis of the statistical velocity gradient’s ellipsoid), direction 2 is the second direction for growth (median axis) and direction 3 is the third direction for growth (minor axis). These directions differed from ROI to ROI. As an example, growth fields and their associated directions at the level of the joint for time window 4—4.5 are shown in Figure 2B(ii). MC and PQ hypertrophic regions were not visible in the cell tracking data, but were included in the FE model of the half-jaw. For these hypertrophic regions, growth rates were set to the average of those of a 30 μm depth of the adjacent proliferative cartilage. In the PQ hypertrophic cartilage, the material orientation of the adjacent proliferative region was used throughout. In the MC hypertrophic region, in which cell orientation varies along the length of the rudiment as shown in Figure 1A, the material orientation of the adjacent cartilage was rotated based on a linear regression of cell orientation with respect to distance from the joint line, fitted to discrete measurements taken in Fiji. The Abaqus user subroutine UEXPAN was used to apply spatially varying expansion based on the strain fields along the corresponding material orientations to provide a prediction of growth and shape for each time-window.

##### Quantification of simulation performance

The predicted shapes were imported into 3-matic where they were aligned with the average jaw shapes of each of the “endpoints” of each time window. Views in the lateral and the ventral planes were exported to Fiji where shape outlines were extracted, and the following shape features were measured: anterior Meckel’s cartilage (MC) length, depth and width and posterior palatoquadrate (PQ) length and depth, as shown in Figure 3. To assess the predictive quality of the simulation for each shape feature, a percentage match of change was calculated as a) the difference between the predicted shape measurement and the initial shape measurement divided by b) the difference between the target shape measurement and the initial shape measurement. The following scores were then assigned based on the percentage match:

- less than 10% match: no growth predicted
- between 10% and 70%: undergrowth
- between 70% and 130%: accurate growth
- above 130%: overgrowth

##### Quantification of the relative roles of growth characteristics

To quantify the relative importance of growth heterogeneity versus growth direction, simulations were conducted in which each of these features was removed or kept constant. Spatial growth heterogeneity was removed in both the MC and PQ by averaging the growth ellipsoids, within the set of ROIs in each rudiment, at each time window. In each rudiment, the average growth ellipsoid was used to generate homogeneous growth maps along the three directions for growth (corresponding to the ellipsoid’s axes) and applied to the model throughout the joint and hypertrophic regions. Orientation in the MC hypertrophic region was still adapted along its length. To remove the role of orientation, isotropic growth was used. Within each ROI in both the joint and hypertrophic regions, an average growth rate corresponding to the average of the three growth ellipsoids’ radii was obtained and applied to the ROI. In other words, ROIs growth ellipsoids became spheres. The resultant shapes when either growth heterogeneity or growth direction were removed were compared to the “full” simulation and with each other.

MATLAB codes for shape averaging, Abaqus CAE models and real and predicted shapes are available at doi: 10.5281/zenodo.5769854.

## Results

### Growth in the zebrafish jaw joint exhibits spatial and temporal heterogeneity as well as marked anisotropy

Comparing shape feature measurements between 3.5 and 5.5 dpf revealed an overall volume expansion over time with a marked increase in Meckel’s cartilage (MC) and palatoquadrate (PQ) length (Figure 3: diamond and triangle), a slight increase in MC and PQ depth (Figure 3: circle and semi-circle) and a slight contraction of MC width (Figure 3: square). In the anterior MC joint element, growth rates in the main direction for growth varied between time windows, ranging from contraction at a mean rate of −2.06 ± 1.49 ×10^−2^ per hour from 3.5—4 dpf, to expansion at a rate of 2.45 ± 0.61 ×10^−2^ per hour from 4.5—5 dpf, as shown in Table 2. In the posterior PQ joint element, growth rates in the main direction consistently increased from a mean rate of 1.01 ± 3.51 ×10^−2^ per hour from 3.5—4 dpf to a mean rate of 2.10 ± 1.27 ×10^−2^ per hour from 5—5.5 dpf, as shown in Table 2. Elevated growth rates in the main direction were observed at the retroarticular process (the most ventroposterior area of the anterior MC joint element shown in Figure 1) from 4—4.5 and 4.5—5 dpf as shown in Figure 4A (black arrows). Growth rates along the second and third directions for growth were much lower than those of the main direction in both the MC and PQ, as shown in Table 2, demonstrating growth anisotropy. Growth maps in the second and third directions for growth are provided in Supplementary Figures 1 and 2. Growth orientations in the anterior MC element exhibited consistent alignment across ROIs; with time, the main direction shifted to align with the ventrodorsal axis from 4.5—5 and 5—5.5 dpf, as shown with solid black lines in Figure 4A. The main direction for growth in the posterior PQ element also tended to align with the ventrodorsal axis from 4.5—5 and 5—5.5 dpf as shown with the black lines in Figure 4B. Overall, growth rates and orientations in the developing jaw joint changed over the time period studied in both joint elements and elevated growth rates were observed at the retroarticular process of the MC demonstrating spatial and temporal growth heterogeneity. Marked growth anisotropy was observed in both joint elements.

**Table 2:**
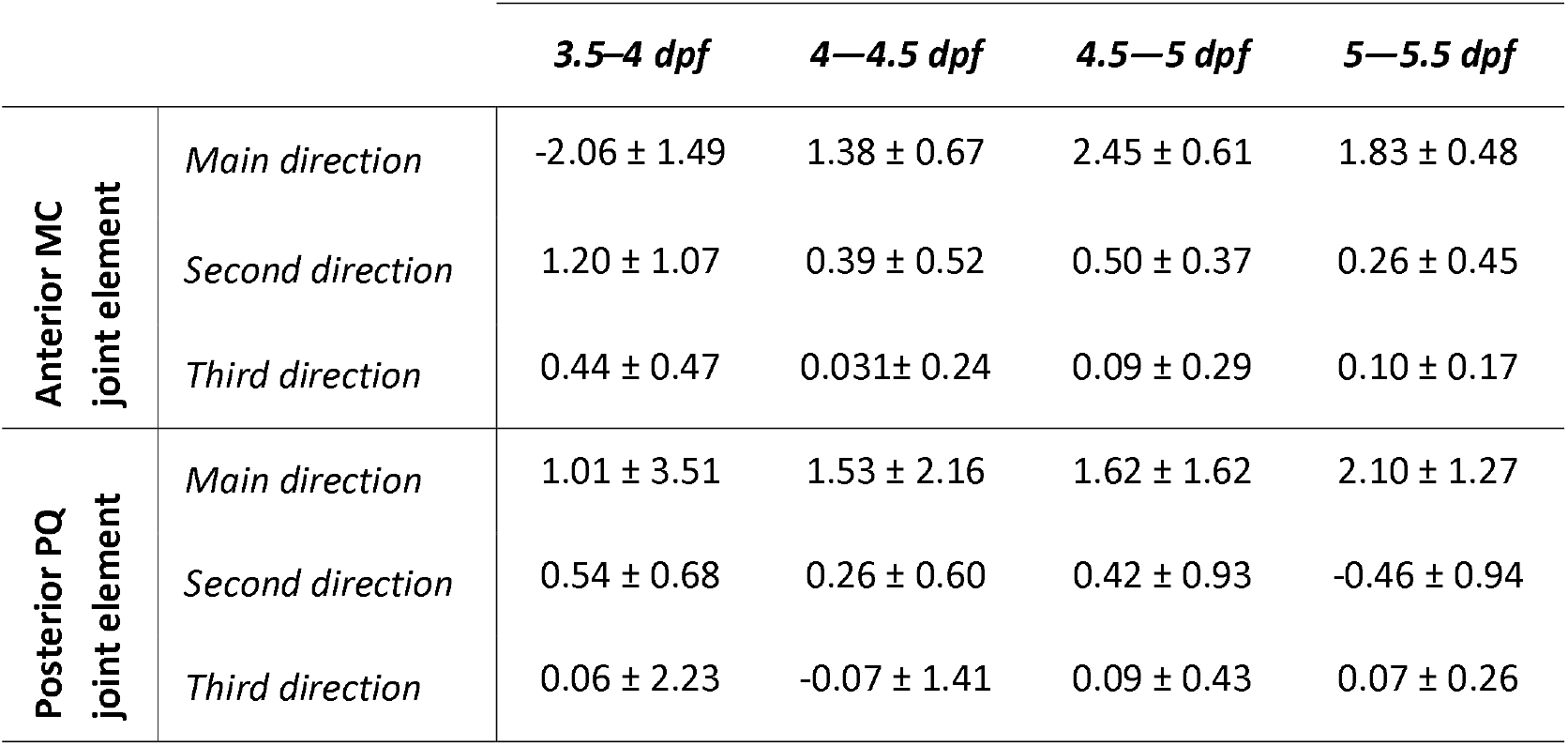
Mean growth rates per hour (×10^−2^) along the three orthogonal directions for growth for each time window (days post fertilization (dpf) 3.5—4, 4—4.5, 4.5—5 and 5—5.5) in the anterior Meckel’s cartilage (MC) and posterior Palatoquadrate (PQ) joint elements.

|  |  | *3.5–4 dpf* | *4–4.5 dpf* | *4.5–5 dpf* | *5–5.5 dpf* |
| --- | --- | --- | --- | --- | --- |
| **Anterior MC**<br>**joint element** | *Main direction* | $-2.06 \pm 1.49$ | $1.38 \pm 0.67$ | $2.45 \pm 0.61$ | $1.83 \pm 0.48$ |
| | *Second direction* | $1.20 \pm 1.07$ | $0.39 \pm 0.52$ | $0.50 \pm 0.37$ | $0.26 \pm 0.45$ |
| | *Third direction* | $0.44 \pm 0.47$ | $0.031 \pm 0.24$ | $0.09 \pm 0.29$ | $0.10 \pm 0.17$ |
| **Posterior PQ**<br>**joint element** | *Main direction* | $1.01 \pm 3.51$ | $1.53 \pm 2.16$ | $1.62 \pm 1.62$ | $2.10 \pm 1.27$ |
| | *Second direction* | $0.54 \pm 0.68$ | $0.26 \pm 0.60$ | $0.42 \pm 0.93$ | $-0.46 \pm 0.94$ |
| | *Third direction* | $0.06 \pm 2.23$ | $-0.07 \pm 1.41$ | $0.09 \pm 0.43$ | $0.07 \pm 0.26$ |

Manual assessment of tracked cells over the time window studied revealed very low proliferation rates in the joint. The percentage of cells which underwent division in the joint over twelve hours was 2.42 ± 1.73 % in the MC and 0.50 ± 0.56 % in the PQ, suggesting that proliferation would only minorly impact on joint growth. No intercalation of joint cells was observed over the twelve-hour timeframes during cell tracking (sample cell tracking over time shown in Figure 5). The volume occupied by tracked joint cells over the timeframe of interest increased substantially, with a mean relative volume expansion per twelve hour period of 18.49 ± 20.44 % in the MC and 23.68 ± 23.92 % in the PQ. Because the ECM forms a thin layer between adjacent cells (see Figure 5), it could not be accurately segmented and its volume was not directly quantified. However, the interstitial space between adjacent cells was consistently narrow, with no apparent increase over time (see example in Figure 5). Therefore, our data indicate that increases in joint volume over the studied time window were primarily due to cell volume increases, rather than proliferation or increases in ECM volume.

**Figure 5:**
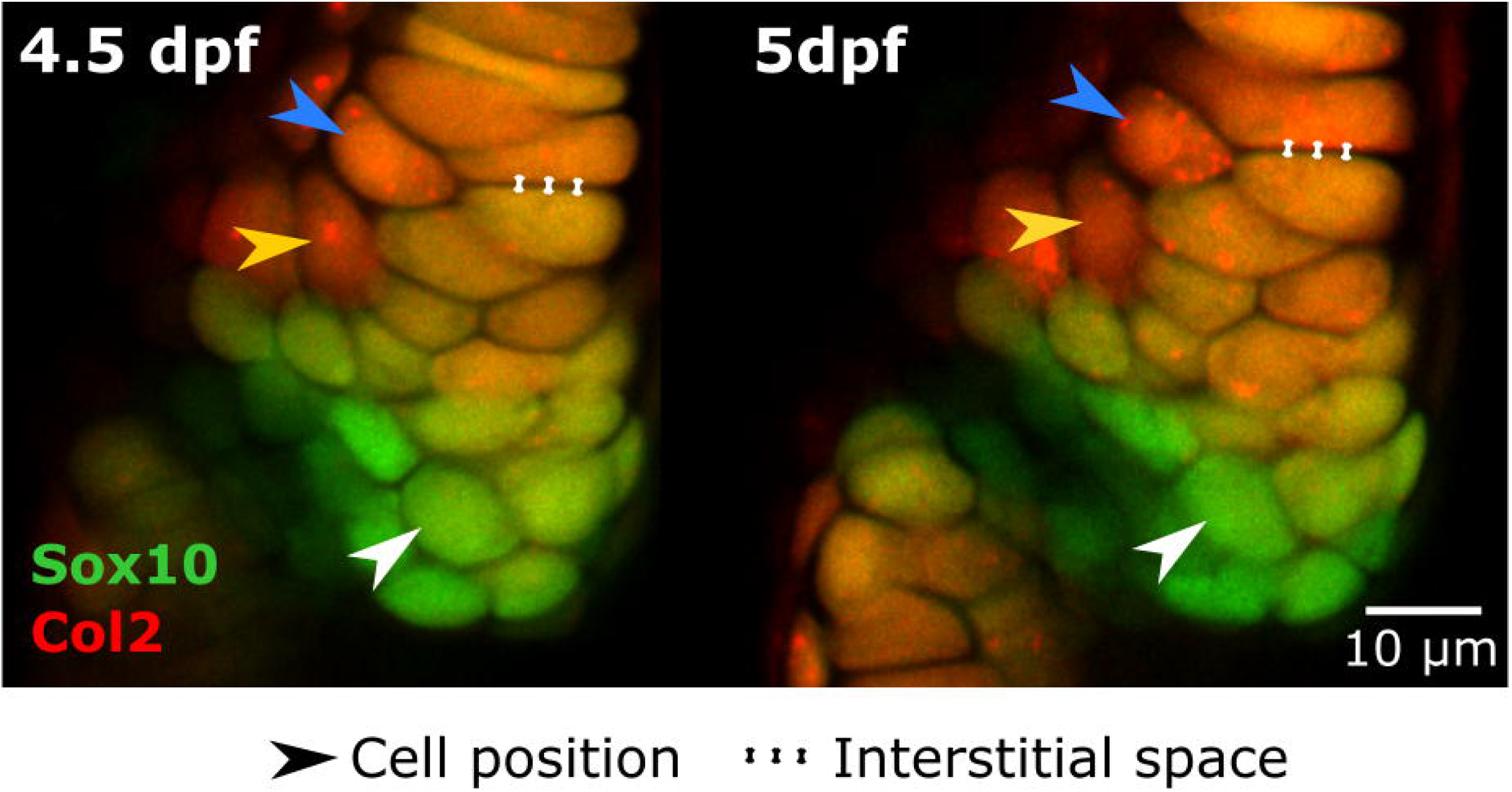
Cell intercalation and the extracellular matrix minorly contribute to jaw joint shaping. Representative ventral stack of the anterior jaw joint element of a live specimen aged 4.5 and 5 dpf expressing the transgenic reporters *Col2a1aBAC:mcherry* and −*4.9sox10:eGFP* marking cartilage. No observation of cell intercalation is made with the cells being clearly identified over time (three cells marked by arrows as examples). The volume occupied by the interstitial space is minor compared to the volume occupied by cells.

### Cell positional information over time enables consistent prediction of zebrafish jaw morphogenesis

Growth for each of the time windows was computationally simulated based on the calculated growth maps, and the shape features undergoing change between 3.5 and 5.5 dpf were used to assess the quality of the shape predictions. For each time window, most observed shape changes were predicted, either accurately, or with some under- or over-growth, as highlighted with green, yellow and purple (respectively) symbols in Figure 6. Length change in both rudiments was accurate from 4—4.5 and 5—5.5 dpf (green triangles and diamonds in Figure 6B & D) but undergrowth was observed from 3.5—4 and 4.5—5 dpf (yellow triangle and diamond in Figure 6A & C). The change of depth in the lateral plane in both rudiments was mostly predicted (yellow and purple circles and semi-circles in Figure 6A, C & D) though only the 4—4.5 predictions accurately matched the target shape (green circle in Figure 6B). The decrease of MC width observed from 3.5—4 and 4.5—5 dpf in the ventral plane was not replicated in the predicted shapes (red squares, Figure 6A & C). Overall, the shape predictions were close to their target shapes (Figure 6). Therefore, zebrafish jaw joint growth and shape change for the time window modelled can be reasonably approximated based on cell positional information over time, where that cell positional information derives mainly from cell rearrangements and volume expansion.

**Figure 6:**
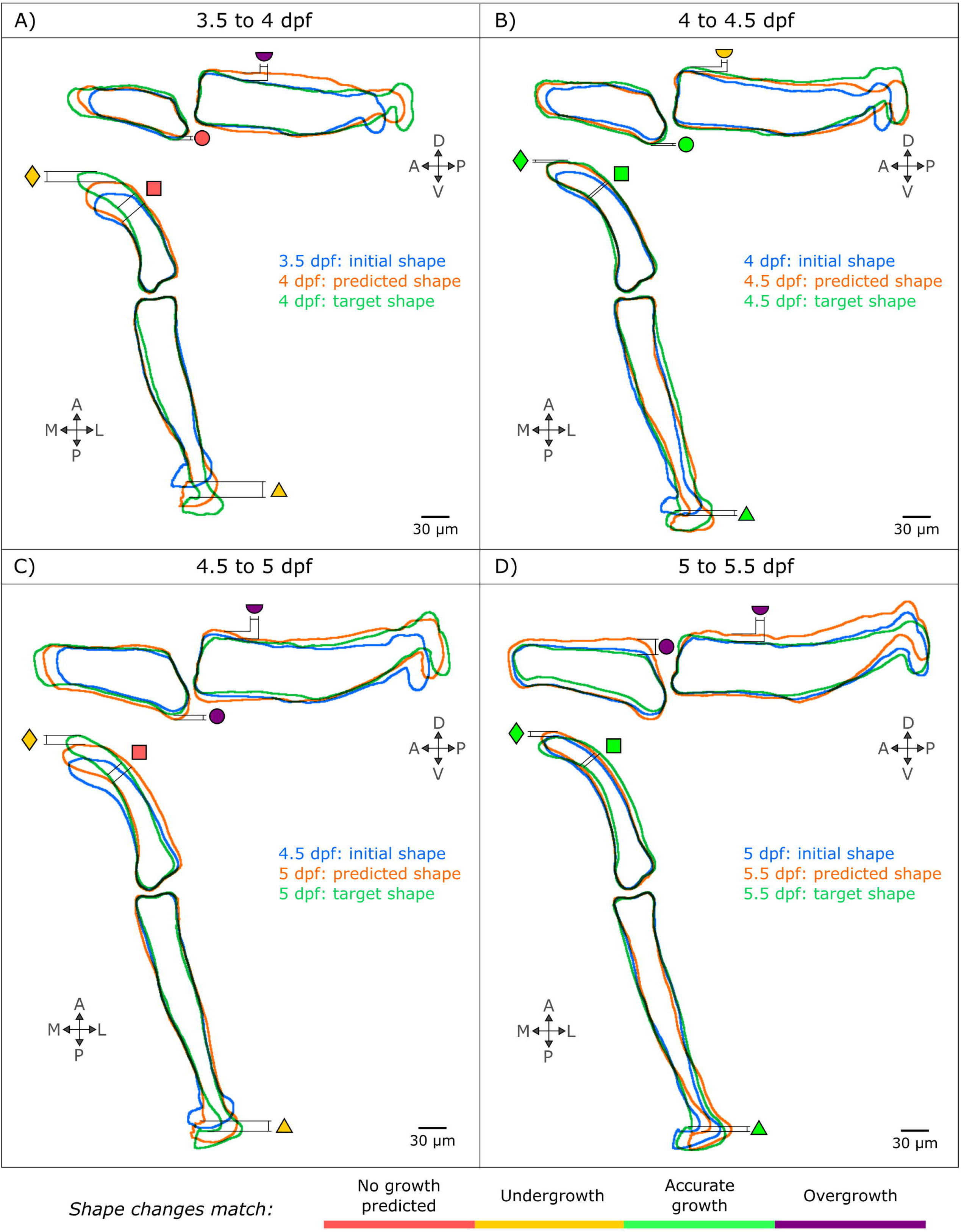
The integration of cell-based data in an FE model successfully predicts zebrafish jaw shape changes from 3.5 to 5.5 dpf, with most faithful predictions from 4 to 4.5 pf. The shape outlines for each time window are superimposed (blue: initial shape, green: target shape, orange: predicted shape) and the shapes features introduced in figure 3 are compared (triangle: Palatoquadrate (PQ) length, diamond: Meckel’s cartilage (MC) length, square: MC width, semi-circle: PQ depth, circle: MC depth) and rated with a colour code explained in the bottom panel (red means no growth predicted, yellow means undergrowth though the pattern of change is correct, green means accurate shape changes and violet means overgrowth though the pattern of change is correct). A: Anterior, P: Posterior, L: Lateral, M: Medial, D: Dorsal, V: Ventral.

### Growth orientation is more important for zebrafish jaw joint shaping than growth heterogeneity

The importance of growth heterogeneity and direction was assessed in simulations in which each of these features was either removed or kept constant. Removing growth heterogeneity resulted in only minor shape changes: over the four time-windows, two features exhibited undergrowth compared to the “full” simulations (PQ length from 3.5—4 dpf and MC depth from 5—5.5 dpf as shown in Figure 7A). In contrast, when growth orientation was removed, several shape features were markedly altered compared to the “full” simulation. From 3.5—4 dpf, both MC and PQ length under isotropic growth exhibited marked undergrowth as seen in Figure 7B. No change was observed from 4—4.5 dpf, while MC depth slightly undergrew from 4.5—5 dpf as shown in Figure 7B. From 5—5.5 dpf, MC and PQ both length and depth were markedly undergrown as shown in Figure 7B. The time windows most severely impacted by the removal of growth orientation (from 3.5—4 and 5—5.5 dpf) were also the windows that exhibited the most complex growth patterns with pronounced growth anisotropy (see Table 2 and Figure 4). Growth predictions for the four time-windows and both adjusted simulation types are provided in Supplementary Figures 3 and 4. These results indicate that growth orientation, and the cellular dynamics likely responsible for it, such as cell orientation and oriented cell division, are crucial to correct morphogenesis. Taken together, our findings suggest that whereas cell proliferation, intercalation and ECM deposition minorly impacted zebrafish jaw joint growth, cell volume expansion and orientation dominate joint growth and morphogenesis.

**Figure 7:**
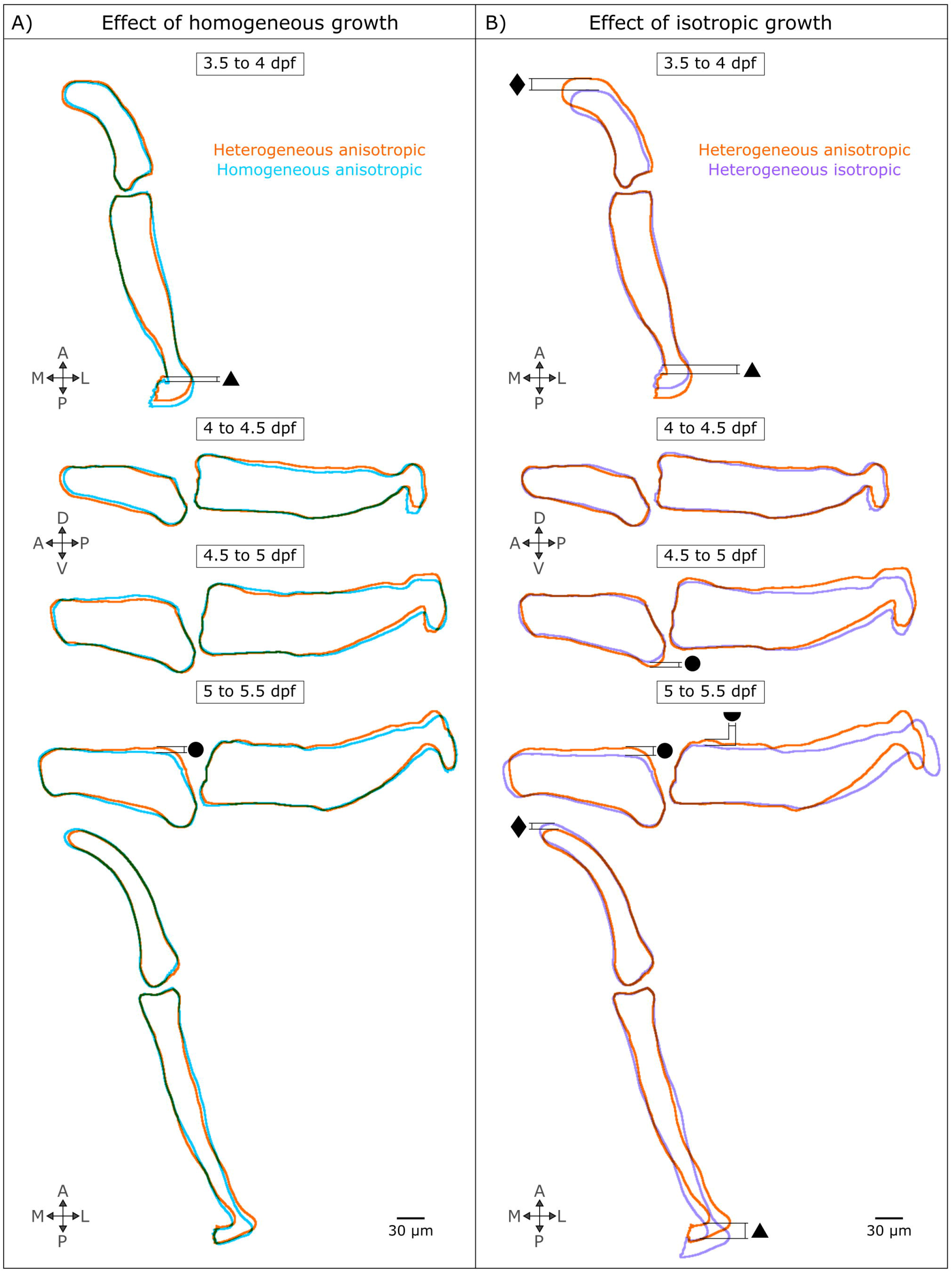
Growth orientation plays an important role in jaw joint shaping whereas growth heterogeneity minorly impacts zebrafish jaw shape predictions. Growth predictions obtained from homogeneous anisotropic (A) and heterogeneous isotropic (B) growth fields are compared with the “full” simulation (heterogeneous anisotropic). Only the views where shape changes were observed in either the homogeneous anisotropic or the heterogeneous isotropic shape predictions are displayed. The shape outlines in all views and time windows are displayed in Supplementary Figures 3 and 4. The black symbols denote the shape features which have been altered when either growth heterogeneity or orientation have been removed (triangle: Palatoquadrate (PQ) length, diamond: Meckel’s cartilage (MC) length, square: MC width, semi-circle: PQ depth, circle: MC depth).

## Discussion

In this research, local tissue deformations of larval zebrafish jaw joints were quantified based upon tracked cell-level data and simulated in a predictive model of joint growth. Our model, the first to simulate joint growth based on biofidelic data, was used to unravel dominant influences and identify which cellular behaviours dominate growth and morphogenesis in the developing zebrafish jaw joint.

Our analysis of zebrafish jaw joint cell dynamics revealed spatially and temporally heterogeneous growth patterns. Growth rates and orientations evolved over the time period studied and elevated growth rates were evident at the retroarticular process of the Meckel’s cartilage, which is known to project ventro-posteriorly from the jaw joint during larval development (Eames et al. 2013). In other developing tissues, such as the developing chick limb bud (Morishita et al. 2015, Suzuki and Morishita 2017) and the drosophilia wing disc (Tozluoglu et al. 2019), spatial and temporal growth heterogeneity was shown to be a key driver of morphogenesis, and in simulations, uniform growth rates did not lead to correct shape predictions (Tozluoglu et al. 2019). In contrast to the limb bud and wing disc, our data indicate that spatial growth heterogeneity is not a dominant influence on zebrafish jaw joint shape for the time windows investigated. Rather, growth orientation was more important for jaw joint growth and morphogenesis in the timeframe studied. Our analysis of zebrafish jaw joint cell dynamics revealed a marked growth anisotropy for the time period studied, and in simulations, isotropic growth led to pronounced shape alterations. This observation is in line with Boehm et al.’s work (2010) in which a parameter optimisation approach on murine limb bud development revealed that growth orientation was critical for accurate shape prediction. Altered cell orientation and increased cell sphericity has been shown to be correlated with altered zebrafish jaw shapes which could indicate a link between cell orientation and growth orientation (Brunt et al. 2015, Brunt et al. 2016, Lawrence et al. 2018).

Our quantification of cell dynamics was derived from cell rearrangements, cell volume expansion and extracellular matrix (ECM) deposition, demonstrating that joint growth and morphogenesis can be reasonably approximated based on these behaviours. Analysis of cell numbers indicated that proliferation is unlikely to be a dominant influence in the joint over the timeframe examined, despite the fact that proliferation has been highlighted in the more mature regions of the developing cartilage elements and in the interzone (Kimmel et al. 1998, Brunt et al. 2017). We also propose that cell intercalation is not likely to have a very strong influence on jaw joint growth in the timeframe and region examined, while acknowledging that cell stacking and convergent extension are key features of more mature regions of the developing cartilage elements (Kimmel et al. 1998, Shwartz et al. 2012, Eames et al. 2013, Mork and Crump 2015, Brunt et al. 2016). As previously reported (Kimmel et al. 1998, Brunt et al. 2016, Brunt et al. 2017), we found that cell volume expansion is likely a key contributor to joint growth, while we found no evidence of substantial increases in ECM volume over the timeframe under investigation. This corroborates the findings of a recent study conducted on the juvenile zebrafish pharyngeal skeleton where ECM volume increase was found to be negligible (Heubel et al. 2021).

Some failures in shape predictions were observed in our results. Cell contraction in the hypertrophic regions of the Meckel’s cartilage has not been accurately simulated due to the specific cell arrangements; in the Meckel’s cartilage, cells stack into a single column in the antero-posterior axis. Because the algorithm for growth quantification does not directly account for cell shape, a medio-lateral contraction of cells in such a columnar arrangement cannot be captured. In addition, under- or over-growth of the Meckel’s cartilage and the palatoquadrate length and depth was observed in some of the timeframes of interest. These imprecisions arise from the small number of cells in the zebrafish jaw. An advantage of our modelling approach will enable it to be applied to organisms with increased numbers of cells and overcome deficiencies resulting from low cell numbers. Because our model focusses on macro-scale shape changes and does not simulate individual cell behaviours, its computational simplicity and practicability enable its use with larger animal models, while cell-based models, such as vertex models in which each cell is represented by a polygon (Alt et al. 2017), are constrained to a limited number of cells.

Our method as presented here is optimal for specimens in which live imaging can be performed. A straightforward application is to quantify growth patterns in epithelial tissues using high cellular resolution images obtained through fluorescence microscopy combined with automated tools for cell segmentation and tracking like EpiTools (Heller et al. 2016). Modelling axolotl joint growth using our approach is also feasible. The axolotl is often used as a model for limb development (Nye et al. 2003, Hutchison et al. 2007) and progress has been made in visualising cells at high resolution during live imaging (Masselink and Tanaka 2021). The existence of rainbow transgenic lines also facilitates cell tracking and visualisation and was used in the past to study digit tip regeneration (Currie et al. 2016). Though live imaging is optimal, it may not be critical to track individual cells with larger numbers of cells. Comparisons between local tissue geometry at successive timepoints may be sufficient to predict joint growth and morphogenesis, which we are exploring in ongoing work.

In conclusion, our findings show that cell volume expansion and orientation are key drivers of larval zebrafish jaw joint growth and morphogenesis. These new insights on what drives joint growth and morphogenesis was facilitated through growth predictions based upon precise and specific cell-level characterisation of growth. Gaining a better understanding of the cell-level processes and dynamics of joint morphogenesis opens up new avenues towards understanding the aetiology of congenital conditions such as developmental dysplasia of the hip and arthrogryposis.

## Acknowledgements

This research was funded by an Anatomical Society PhD studentship to J.G.. C.L.H. was funded by Versus Arthritis Fellowship 29137. E.L. was funded by a Wellcome Trust Dynamic Cell PhD studentship. M.W. was funded by the China Scholarship Council. We thank James Monsen for providing the methodology and MATLAB script which was used for generating average shapes. We would like to thank Mat Green for zebrafish husbandry and the staff of the Wolfson Bioimaging centre Bristol for imaging support.

## Author contributions

Conceptualisation: J.G., C.L.H., N.C.N.; Methodology: J.G., E.A.L., M.W., C.L.H., N.C.N.; Software: J.G.; Analysis and visualisation: J.G., C.L.H., N.C.N.; Supervision: C.L.H., N.C.N.; Writing – original draft: J.G., N.C.N.; Writing – review & editing: J.G., E.A.L., M.W., C.L.H., N.C.N.; Funding acquisition: C.L.H., N.C.N.

## Supplementary information

Data underlying this article can be accessed on zenodo at doi: 10.5281/zenodo.5769854.

