## Supplemental Data 1 for "Growth orientations, rather than heterogeneous growth rates, dominate jaw joint morphogenesis in the larval zebrafish"

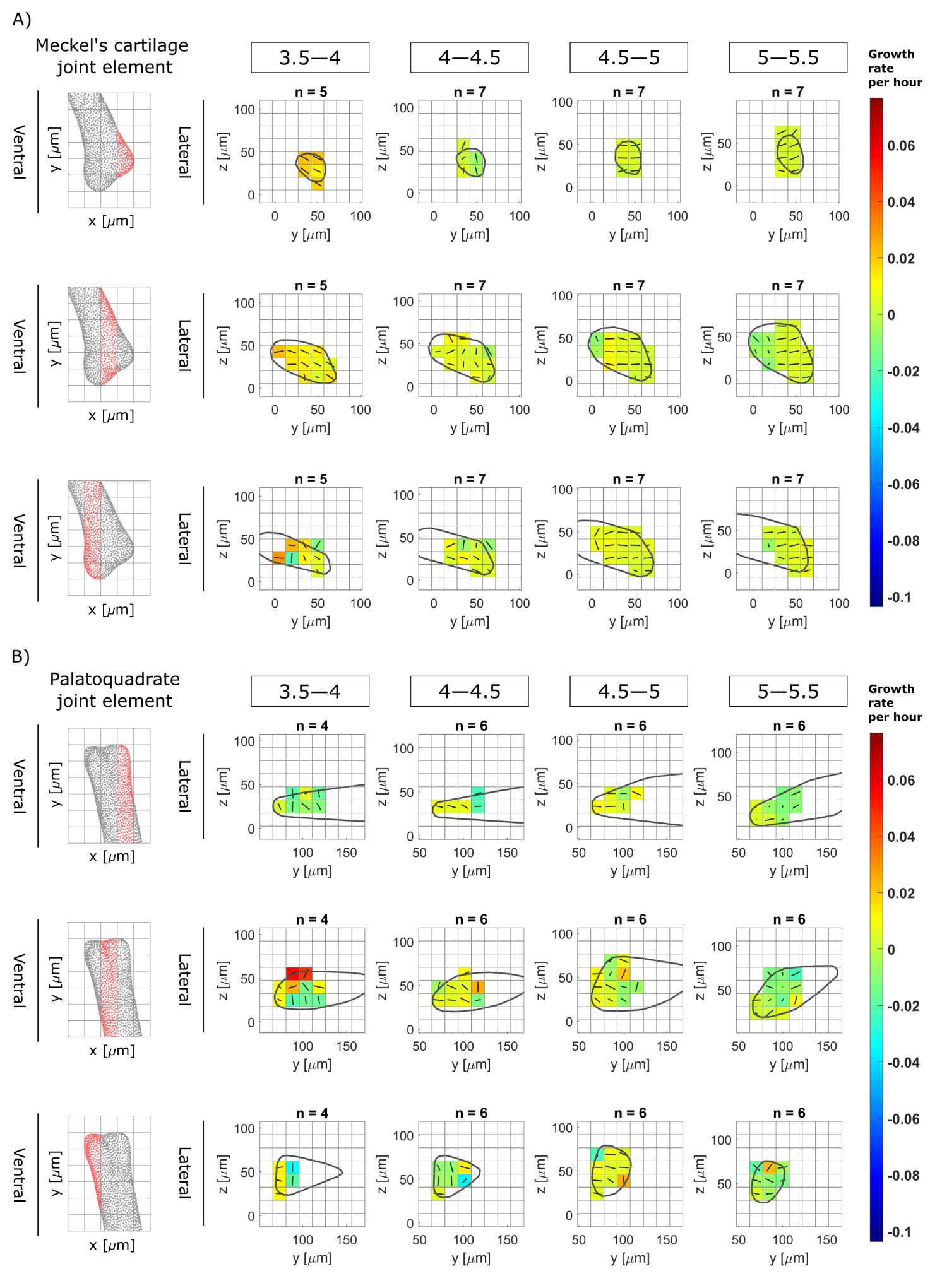

**Supplementary figure 1:** Maps showing growth rates along the second direction for growth (median axis of the ellipsoid) and their associated directions for each time window (3.5—4, 4—4.5, 4.5—5 and 5—5.5) in the anterior Meckel’s cartilage (A) and posterior Palatoquadrate (B) joint elements in the lateral plane. Growth rates are represented by colours while the direction is shown by solid black lines. Results are displayed across the rudiment’s depth; views in the ventral plane of each section are displayed on the left panels.

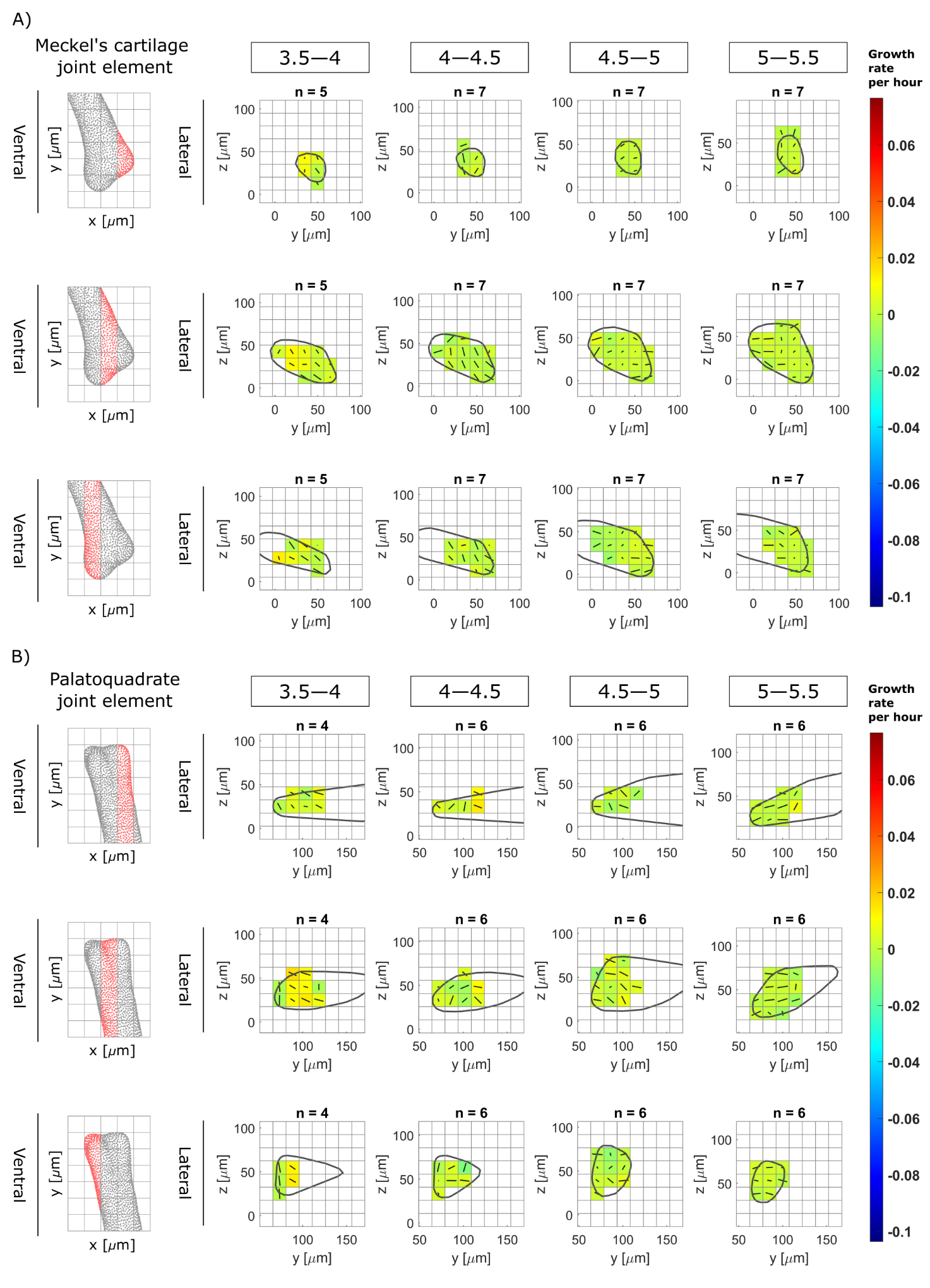

**Supplementary figure 2:** Maps showing growth rates along the third direction for growth (minor axis of the ellipsoid) and their associated directions for each time window (3.5—4, 4—4.5, 4.5—5 and 5—5.5) in the anterior Meckel’s cartilage (A) and posterior Palatoquadrate (B) joint elements in the lateral plane. Growth rates are represented by colours while the direction is shown by solid black lines. Results are displayed across the rudiment’s depth; views in the ventral plane of each section are displayed on the left panels.

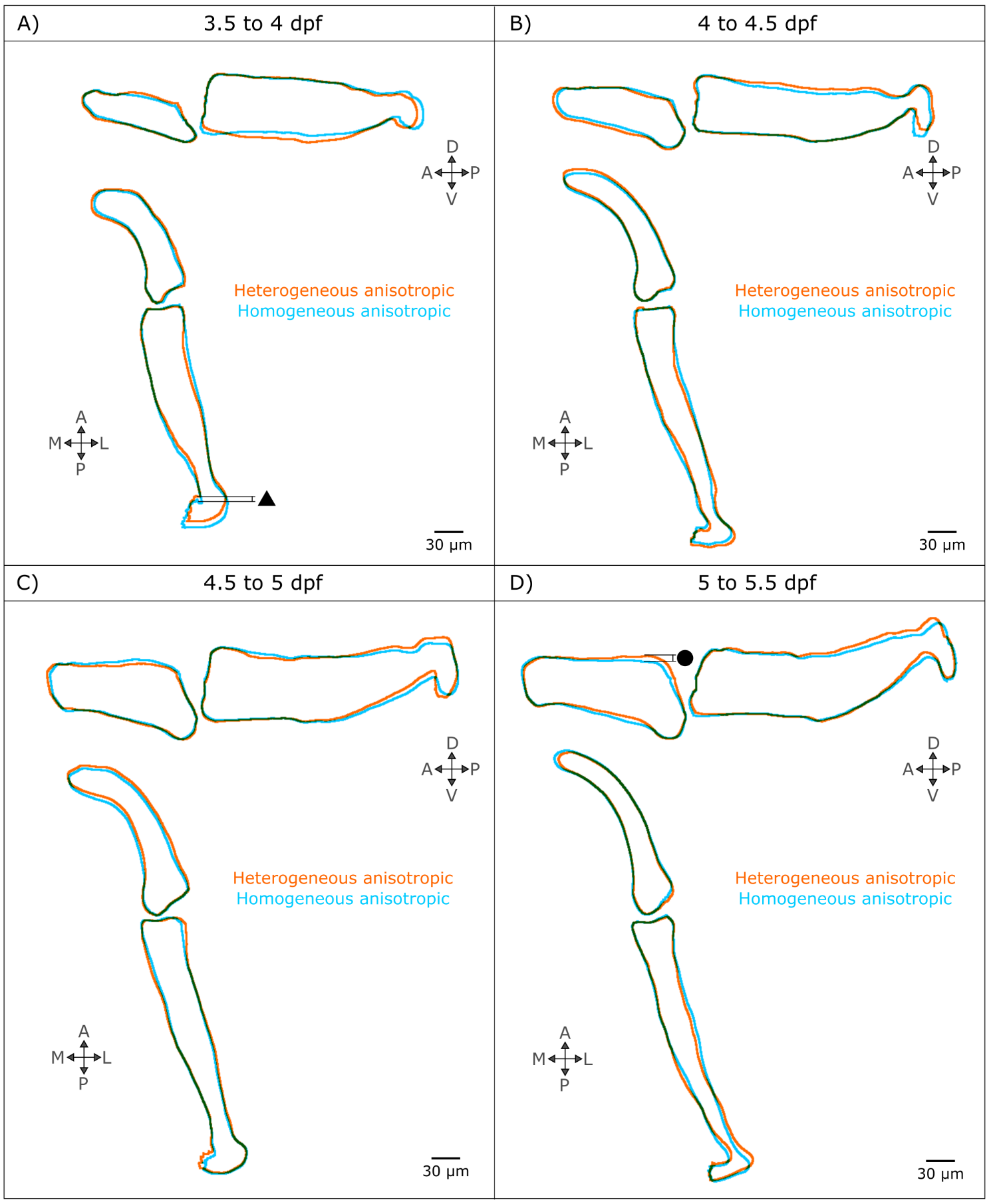

**Supplementary figure 3:** Growth predictions obtained from homogeneous anisotropic growth fields are compared with the “full” simulation for each time window. The black symbols denote the shape features which have been altered when growth heterogeneity has been removed (triangle: Palatoquadrate (PQ) length, diamond: Meckel’s cartilage (MC) length, square: MC width, semi-circle: PQ depth, circle: MC depth).

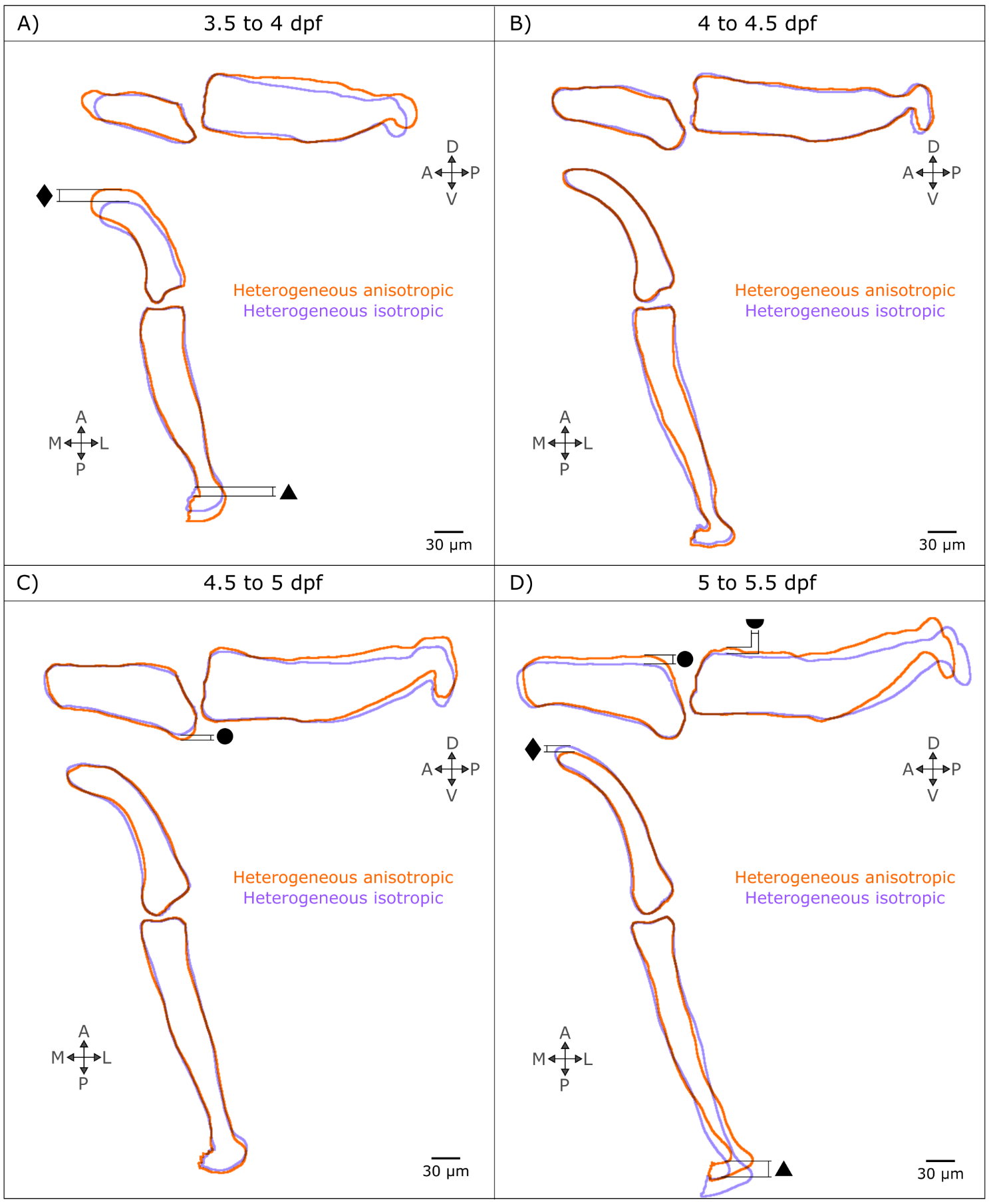

**Supplementary figure 4:** Growth predictions obtained from heterogeneous isotropic growth fields are compared with the “full” simulation for each time window. The black symbols denote the shape features which have been altered when growth orientation has been removed (triangle: Palatoquadrate (PQ) length, diamond: Meckel’s cartilage (MC) length, square: MC width, semi-circle: PQ depth, circle: MC depth)
